# Sialoglycans on human T cells attenuate death programs executed through the Fas pathway

**DOI:** 10.1101/2025.09.15.676380

**Authors:** Vanessa Affe, Qianmeng Lei, Tim S. Veth, Emmajay Sutherland, Hani Choksi, Fauzia N. Izzati, Qingyu Shi, Haissi Cui, Nicholas M. Riley, Landon J. Edgar

## Abstract

T cells are critical executors of adaptive immune responses and their persistence is tightly regulated. Part of this regulation relies on programmed cell death driven by the Tumor Necrosis Factor (TNF) receptor superfamily. The addition of glycans that terminate in the monosaccharide sialic acid (sialoglycans) to these cell death receptors has been shown to attenuate their apoptotic functions. While this is now understood to be a pro-survival mechanism in settings of cancer pathophysiology, the specific roles of sialoglycans in regulating cell death receptor activity on human T cells remains unexplored. This is of particular importance given the rising interest in T cell glycan editing for therapeutic benefit. Here, we address this gap using both immortalized (Jurkat) and primary human T cells deficient in sialoglycans. We found that T cell sialoglycans suppressed apoptosis induced by the Fas receptor (FasR) but not other TNF receptor superfamily members such as TNFR1 and TRAIL-R1. Dynamic reorganization of FasR was increased on sialoglycan-deficient Jurkat cells, suggesting that these glycans limit receptor clustering. This model was further supported by phosphoproteomics results, which confirmed that loss of sialoglycans negatively regulated the pro-survival MAPK/ERK signalling pathway. Finally, we used a recombinant sialic acid cleaving enzyme (sialidase) to confirm that sialoglycans on primary human T cells are *bona fide* immunophysiological regulators of FasR-driven programmed cell death. Combined, our results demonstrate that sialoglycan remodelling on T cells influences cell fate driven by the Fas pathway and provide motivation to further characterize the immunoregulatory roles of the glycocalyx in health and disease.

## Introduction

Cell death receptors of the tumor necrosis factor (TNF) superfamily play critical roles in regulating leukocyte homeostasis (1). The best studied members of this superfamily include TNF receptor 1 (TNFR1), the FS-7-associated surface antigen (FasR/TNFRSF6) (2), and TNF-related apoptosis-inducing ligand (TRAIL-RI/TNFSF10A). While all of these transmembrane proteins share substantial sequence homology in their extracellular domains (3), sufficient diversity exists to enable selective recognition of various death receptor ligands. Engagement with a cognate ligand – TNF-α for TNFR1, FasL for FasR, and TRAIL for TRAIL-RI – initiates cytotoxic signals by promoting receptor aggregation and recruitment of adaptor proteins to the conserved intracellular death domain (4). Downstream phosphorylation events ultimately activate select caspases which can drive cell death programs (5). Signalling through the death domain can also regulate non-apoptotic events including inflammation, cell differentiation, and pro-tumourogenic programs (4, 5).

While ligand engagement is the key event that initiates signalling through the various death receptors, other factors can influence the nature/intensity of the resulting response. For example, TNF superfamily receptors – like most proteins on the cell surface – are glycosylated (6, 7), and the specific composition of these glycans is known to add an additional level of regulation to these pathways. Specifically, a series of key studies by Bellis and coworkers (6, 8–11) have shown that N-linked glycans terminating in sialic acid residues (sialoglycans) can attenuate clustering and internalization of TNFR1 and FasR, which ultimately decreases cell death in response to TNF-α and FasL respectively. These effects were uniquely driven by the sialoglycan products produced by the sialyltransferase enzyme ST6GAL1, which attaches sialic acid in an α2-6-configuration to galactose on N-linked glycans (12). Most of this work was executed using cell lines derived from various cancers, many of which can increase sialyltransferase as a pro-survival mechanism (13, 14). While these previous studies have clearly demonstrated a functional connection between α2-6-sialoglycan production and death receptor activity in the context of cancer growth and survival, the relevance of these observations to non-pathophysiological systems has been less explored. One study used a differentiated cell line and murine models to show that upregulation of ST6GAL1 prevents TNF-α-promoted apoptosis of macrophages (6). The effects of death receptor sialylation on leukocyte immunophysiology beyond this example remain unexplored.

We and others have demonstrated that removing sialic acids from the surface of T cells – critical mediators of adaptive immunity – influences their phenotype and functions (15–20). There is now significant interest in leveraging technologies that decrease sialic acid density on T cells to potentiate their activity in settings of functional exhaustion (16, 18, 20). While this approach has important potential therapeutic applications, it is possible that broad desialylation of a T cell surface could also potentiate sensitivity to death ligand-promoted cell death. This would be undesirable in settings where the goal is to promote an enhanced T cell response but may be of benefit in scenarios involving T cell driven autoimmunity. Here, we explore this possibility by evaluating the impact of decreased sialylation on TNF receptor superfamily programmed cell death in human T cells. We established a model T cell system that lacked the capacity to produce α2-6-sialoglycans (*ST6GAL1*^-/-^) and found that these cells were more susceptible to FasL-triggered apoptosis. Imaging flow cytometry revealed that reorganization of FasR was more dynamic on *ST6GAL1*^-/-^ cells following treatment with FasL, consistent with previous reports of enhanced mobility of TNFR1 lacking α2-6-sialylation. We found that one consequence of this altered FasR mobility was a dramatic change in intracellular phosphorylation events, including those known to be involved in FasR signalling. Finally, we confirmed that global removal of sialic acids from primary human T cells increased their susceptibility to FasL-promoted apoptosis. This increase was maximized on stimulated T cells, which are known to display enhanced α2-6-sialylation compared to naïve counterparts (19). Taken together, these results confirm that sialoglycans regulate human T cell sensitivity to FasR signalling. They also underscore the potential complexities associated with manipulating glycans on T cells for therapeutic benefit.

## Results

### A model T cell system deficient in α2-6-sialoglycans

Jurkat cells are an immortalized T cell line originally derived from the peripheral blood of a male leukemia patient (21). They have been used extensively to study human T cell immunophysiology as they are trivial to culture and compatible with gene editing technologies. We measured substantial α2-6-sialoglycan presentation on wild-type (WT) Jurkat cells via staining with *sambucus nigra* lectin (SNA) in a flow cytometry assay (Fig. 1, *A* and *B*). Quantitative real-time PCR (RT-qPCR) revealed that the cells expressed ST6GAL1 (Fig. S1*A*), which is known to manufacture the α2-6-sialoglycans recognized by SNA (12). Transcript levels for *ST6GAL1* were even higher than the housekeeping gene *GAPDH* when normalized to 18S ribosomal RNA, suggesting a very high capacity for production of α2-6-sialoglycans in these cells, much like activated primary human T cells (19). We next confirmed that Jurkat cells expressed FasR, modest amounts of TNFR1, and low-to-negligible levels of TRAIL-RI using flow cytometry (Fig. 1, *C* and *D*). Combined, these data revealed that Jurkat cells possessed the critical components required to study the roles of α2-6-sialoglycans on cell death programs executed by the TNF receptor superfamily.

**Figure 1.**
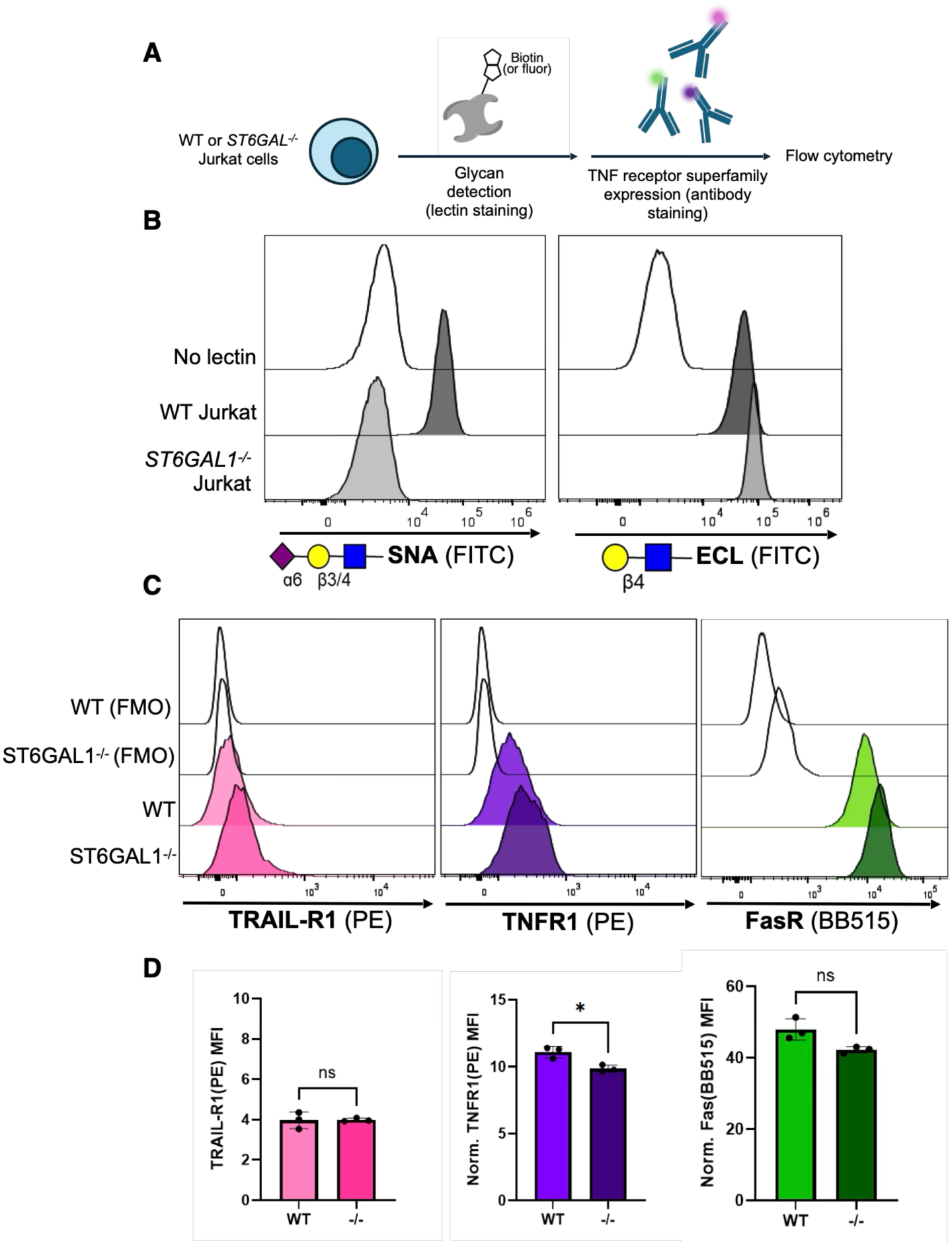
Loss of biosynthetic capacity for α2-6-sialoglycans minimally influences cell death receptor expression on a model T cell line. *A*, ST6GAL1 knock out Jurkat cells were generated using CRISPR/Cas9 and phenotyped via flow cytometry. *B*, Presentation of α2-6-sialoglycans and their corresponding asialo-*N*-acetyllactosamine counterparts was measured on WT and *ST6GAL1^-/-^* Jurkat cells using the fluorescent lectins from *Sambucus Nigra* (SNA) and *Erythrina Cristagalli* (ECL), respectively. C, Expression of the TNF receptor superfamily members TRAIL-R1, TNFR1, and FasR on WT and *ST6GAL1^-/-^*Jurkat cells. *D*, Quantification of data from *C*. Results are reported as mean SD from three experiments. \**p* < 0.05, ns = *p* > 0.05 = not significant, student’s t-test, two-tailed.

To generate Jurkat cells without α2-6-sialoglycans, we used CRISPR/Cas9 to knock out (KO) the gene coding for ST6GAL1. Staining of the *ST6GAL1*^-/-^ cells with SNA produced the same signal as the background staining control (Fig. 1*B*), consistent with our expectation that the edit would abolish all α2-6-sialoglycan biosynthesis in these cells. We determined that loss of *ST6GAL1* only minimally impacted expression of TNFR1 and did not significantly impact levels of FasR or TRAL-RI (Fig. 1, *C* and *D*).

### Loss of α2-6-sialylation increases susceptibility to FasL-induced apoptosis in Jurkat cells

We next used the WT and *ST6GAL1^-/-^* Jurkat cells to determine if loss of α2-6-sialylation impacted cell death programs executed through the TNF receptor superfamily. Both cell lines were treated with TNF-α, FasL, and TRAIL (Fig. 2*A*). After incubation, we stained the cells with Annexin V and DAPI to identify live, apoptotic and dead populations using flow cytometry (Fig. 2*B*). We specifically focused on cells from the live and apoptotic populations as we were interested in overall viability and early programmed cell death as a function of sialylation and cognate ligand binding. A small increase in apoptosis following exposure to TRAIL was detected, with more *ST6GAL1*^-/-^ cells impacted compared to WT counterparts (Fig. 2*C*). A similar effect was observed for cells treated with TNF-α (Fig. 2*D*). In contrast, treatment with FasL induced a dramatic increase in apoptosis uniquely in the *ST6GAL1*^-/-^ cells, whereas WT counterparts were impacted similarly to the other death ligands (Fig. 2*E*). A dose-response analysis confirmed the effect was magnified with increasing amounts of FasL (Fig. S2).

**Figure 2.**
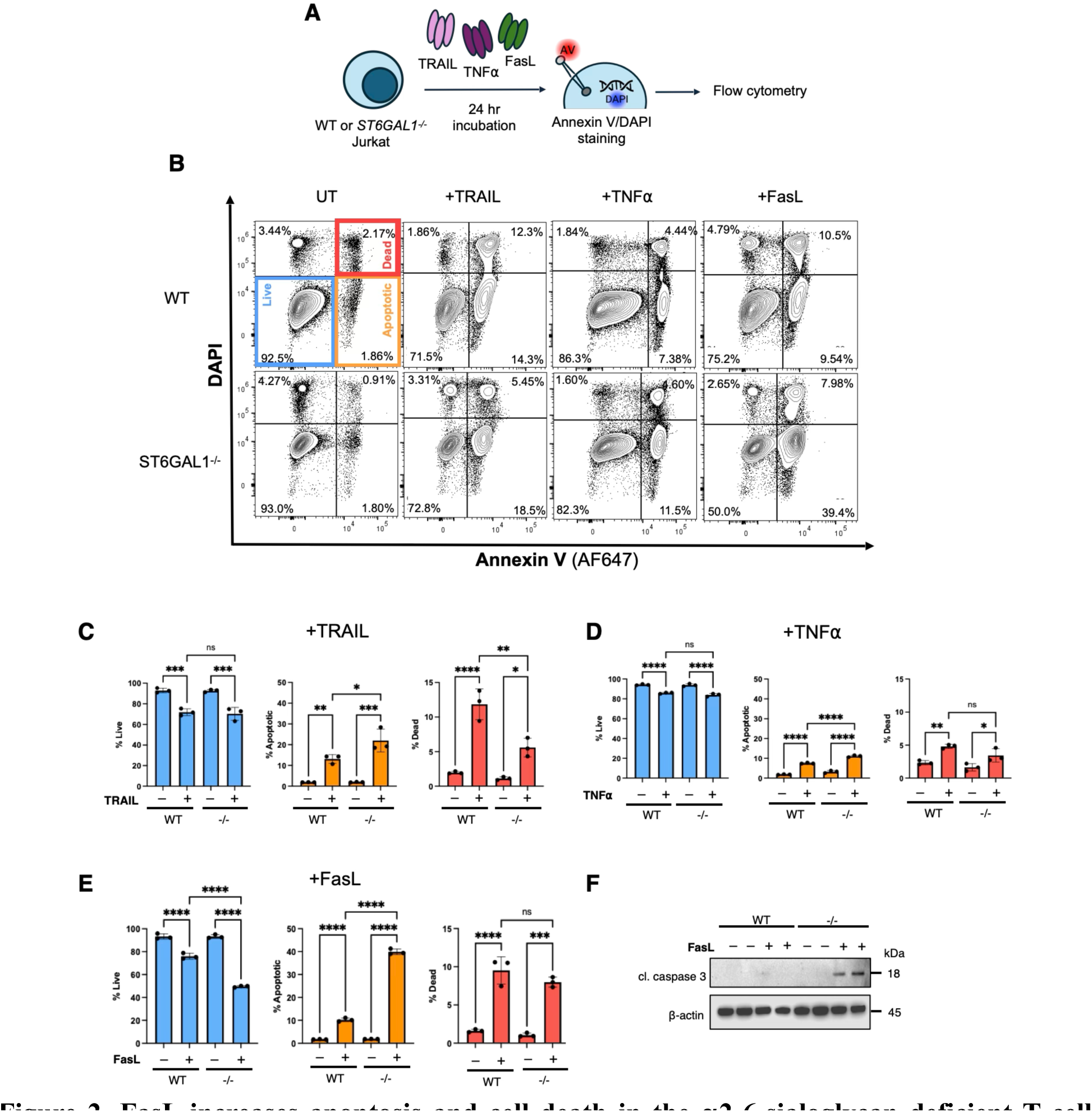
FasL increases apoptosis and cell death in the α2-6-sialoglycan deficient T cell model compared to other TNF receptor superfamily ligands. *A*, Workflow for evaluating WT or *ST6GAL1^-/-^*Jurkat cell death induced by TRAIL, TNF-α, and FasL. Annexin V (recognizes exposed phosphatidylserine) and DAPI (intercalates DNA) report on apoptosis and cell death, respectively. *B*, Flow cytometry plots for Annexin V and DAPI staining of WT or *ST6GAL1^-/-^*Jurkat cells treated with 100 ng/mL TRAIL, TNF-α, and FasL. Bottom left gates (blue) = live cells, bottom right gates (orange) = apoptotic cells, and top right gate (red) = dead cells. Percent values indicate the portion of total singlet cell events within each quadrant. *C*, *D*, and *E*, Quantification of data from *B*. *F*, Immunoblot against cleaved (cl.) caspase 3 (18 kDa) with β-actin shown as loading control (45kDa). For plots in *C*, results are reported as mean SD from three experiments. \**p* < 0.05, \*\**p* < 0.01, \*\*\**p* < 0.001 \*\*\*\**p* < 0.0001, *p* > 0.05 = ns = not significant. Two-way ANOVA with Tukey post-hoc test.

To test if this effect was associated with the FasR signalling pathway, we evaluated cleaved caspase 3 (CC3) levels in WT and *ST6GAL1*^-/-^ cells treated with FasL compared to untreated controls. Production of CC3 is stimulated following FasL engagement with FasR and is required for the death inducing signalling complex in lipid rafts (22). Immunoblotting revealed that CC3 levels were greatly elevated in *ST6GAL1*^-/-^ cells and below the limit of detection in the WT group (Figs. 2*F* and S3). These results provided evidence that the increased apoptotic response in *ST6GAL1^-/-^* Jurkat cells upon exposure to FasL was likely due to enhanced signalling through the death domain of FasR.

### FasR reorganization is more dynamic on α2-6-sialoglycan deficient Jurkat cells

With the knowledge that *ST6GAL1*^-/-^ cells were more sensitive to FasL induced apoptosis, we next wanted to understand if spatial reorganization and/or internalization of FasR was a potential driver of the effect. We were motivated to study these parameters since others have reported that TNFR1 oligomerization and internalization was potentiated on *ST6GAL1*^-/-^ HEK293 cells following treatment with TNF-α (11). Here, we leveraged imaging flow cytometry (23) to measure FasR expression and organization on live, apoptotic, and dead cells (Fig. 3*A*) using the same Annexin V/DAPI approach as in Fig. 2*B*. These data revealed that absolute FasR signal (MFI) was diminished only on apoptotic *ST6GAL1^-/-^* cells treated with FasL compared to the corresponding live population (Fig. 3*B* and *C*). While we detected moderately higher surface presentation of FasR on *ST6GAL1*^-/-^ cells, the higher autofluorescence of the KO line likely contributed to this apparent increase (Fig. 1*C*). This assay did not detect FasL-dependent differences in surface presentation of FasR across WT groups under these conditions. Taken together, these results suggested that more dramatic internalization of FasR on *ST6GAL1^-/-^* cells occurred compared to WT companions, which was consistent with previous studies of FasR internalization in *ST6GAL1^-/-^* HD3 cells (colorectal cancer) (8).

**Figure 3.**
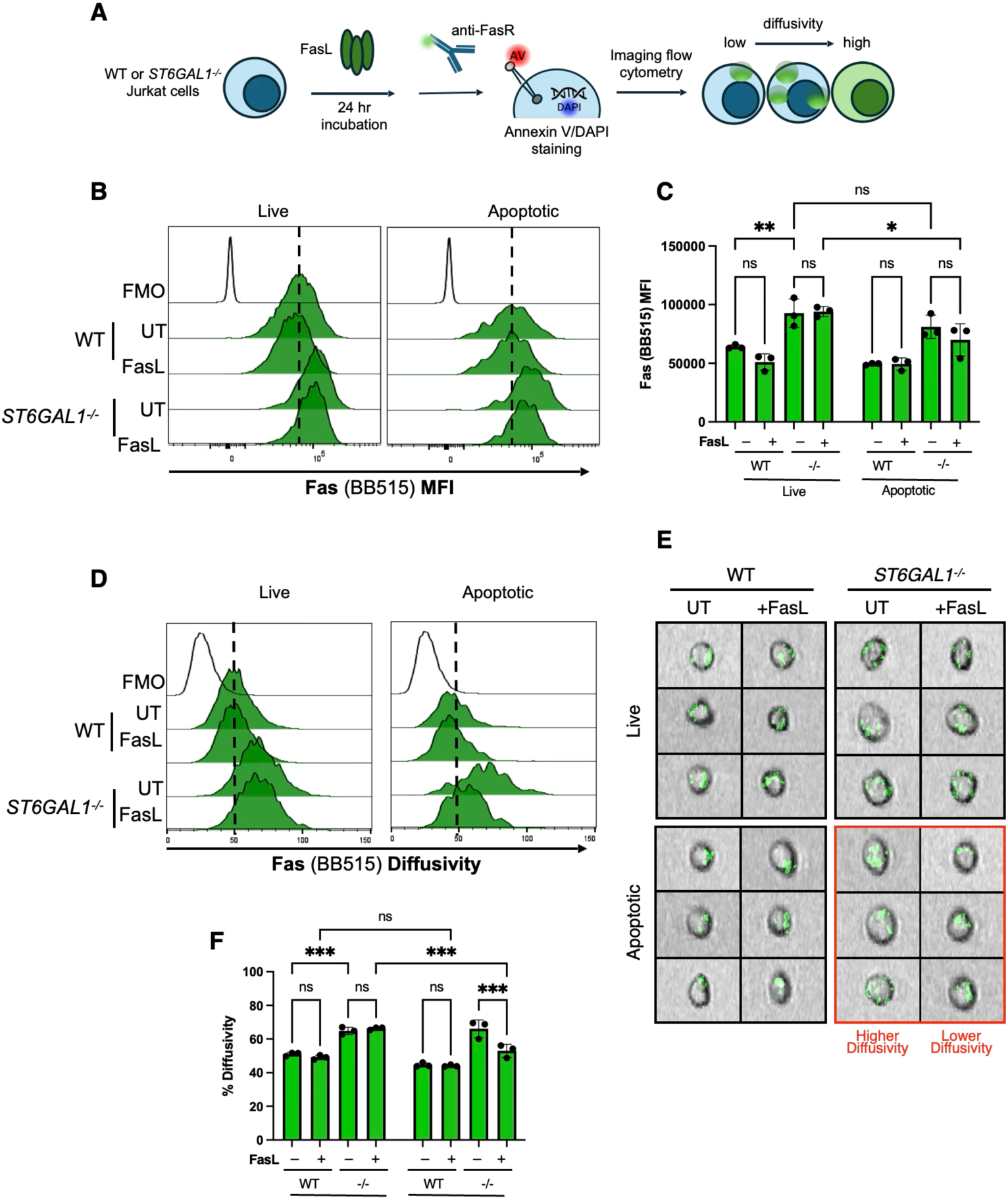
a2-6-sialoglycan deficiency enhances localization dynamics of the Fas Receptor. *A*, Workflow for evaluating surface presentation and spatial organization of FasR on WT or *ST6GAL1^-/-^* Jurkat cells via imaging flow cytometry. *B*, Surface expression levels of FasR as a function of ST6GAL1 expression, exposure to FasL (100 ng/mL), and live vs. apoptotic status as measured by the BB515 channel on the imaging flow cytometer (same dataset as imaging data in subsequent panels). Gating scheme for live vs. apoptotic as in Fig. 2B. FMO = fluorescent minus one staining control. *C*, Quantification of results from *B*. *D*, Representative individual cell images from the same groups of cells as reported in *B*. Data recorded via imaging flow cytometry. Greyscale layer corresponds to lightloss, which is analogous to a brightfield image of the cell. Green layer corresponds to the fluorescence signal from the anti-FasR antibody. *E*, Diffusivity of signal from the fluorescent anti-FasR antibody. *F*, Quantification of data from *E*. For plots in *C* and *F*, results are reported as mean SD from three experiments. \**p* < 0.05, \*\*\**p* < 0.001, \*\*\*\**p* < 0.0001, *p* > 0.05 = ns = not significant. Two-way ANOVA with Tukey post-hoc test.

We next evaluated the diffusivity of FasR signals on individual cell images as a metric for FasR organization (23). Low diffusivity values indicate that FasR is localized in discrete clusters whereas high values suggest less organization into clustered units. We found that live WT cells had lower FasR diffusivity compared to the corresponding *ST6GAL1*^-/-^ cells (Fig. 3*D*, *E*, and *F*). Diffusivity was unchanged in live cells from both groups upon treatment with FasL, suggesting that FasR reorganization could not be observed at this timepoint or until cells entered an apoptotic program. In contrast, we found that FasR diffusivity on apoptotic *ST6GAL1*^-/-^ cells was significantly decreased upon treatment with FasL (Fig. 3*F*). We did not detect changes in diffusivity of FasR on WT cells between live and apoptotic populations which suggested that the extent of reorganization of FasR in settings of complete α2-6-sialylation was beyond the limit of detection of this imaging flow cytometry assay. Combined, these data suggest that FasL-promoted reorganization of FasR was more dynamic on cells deficient in α2-6-sialoglycans. This partially explained why *ST6GAL1^-/-^* cells were more suspectable to FasL-induced apoptosis since clustering of FasR is required for initiation of cell death programs.

### Loss of α2-6-sialoglycans negatively regulates the MAPK/ERK signalling pathway in response to FasL

While surface organization and internalization of FasR are important parameters related to initiation of signalling, these metrics do not report on the downstream intracellular phosphorylation events that are central to cell death programs (24). To probe this, we leveraged a mass spectrometry-based phosphoproteomics workflow to measure relative changes in Ser/Thr phosphosite occupancy on intracellular proteins following treatment with FasL (Fig. 4*A*). This approach enabled site-specific quantification across the human phosphoproteome.(25, 26) We detected many proteins with differential phosphosite occupancy between WT and *ST6GAL1^-/-^* cells treated with FasL (Fig. 4*B*). This confirmed that a relationship existed between loss of α2-6-sialylation and intracellular phosphorylation events. The differences included both increased and decreased phosphosite occupancy across a range of proteins, suggesting that both kinase and phosphatase activity was impacted by the loss of α2-6-sialylation. We used the Robust Inference of Kinase Activity application (RoKAI) designed by Yilmaz and coworkers (27) to identify the specific kinases and phosphatases that were responsible for the significantly changed phosphosite occupancies (Figs. 4*C* and S4). Here, we found that several kinases and phosphatases known to be involved in FasR signalling were differently regulated – especially those from the MAPK/ERK pathway (28–30) – across *ST6GAL1^-/-^* and WT Jurkat cells treated with FasL. Specifically, *ST6GAL1^-/-^* cells exhibited decreased ERK 1/2 activity and increased dephosphorylation executed by protein-tyrosine phosphatase non-receptor type 7 (PTPN7) – an ERK-specific phosphatase (31, 32). We also detected increased activity of several dual-specific phosphatases (DUSP1/2/3), which are known to dephosphorylate the MAPK family of proteins (31, 32). In addition to this, the activity of MEK1/2 – an upstream kinase in the MAPK/ERK signalling pathway (33, 34) – was also decreased in the FasL-treated *ST6GAL1^-/-^* cells. These changes in kinase and phosphatase activity were associated with the cell fate-related gene ontology (GO) biological process terms: ‘apoptotic processes’ (GO:0006915), ‘programmed cell death’ (GO:0012501), ‘regulation of apoptotic processes’ (GO:0042981), and ‘regulation of programmed cell death’ (GO:0043067) (Fig. 4*D*). While significant differences in phosphopeptide abundance were detected in *ST6GAL1^-/-^* vs. WT cells even without FasL treatment (Fig. S4), only two kinases could be assigned with statistically significantly changes in activity through RoKAI analysis – PKCA and PKCE. Together, these results suggested that attenuated phosphorylation through MAPK/ERK in parallel with increased DUSP-catalyzed dephosphorylation was a dominant mechanism through which loss of α2-6-sialoglycans potentiated FasL-induced cell death programs.

**Figure 4.**
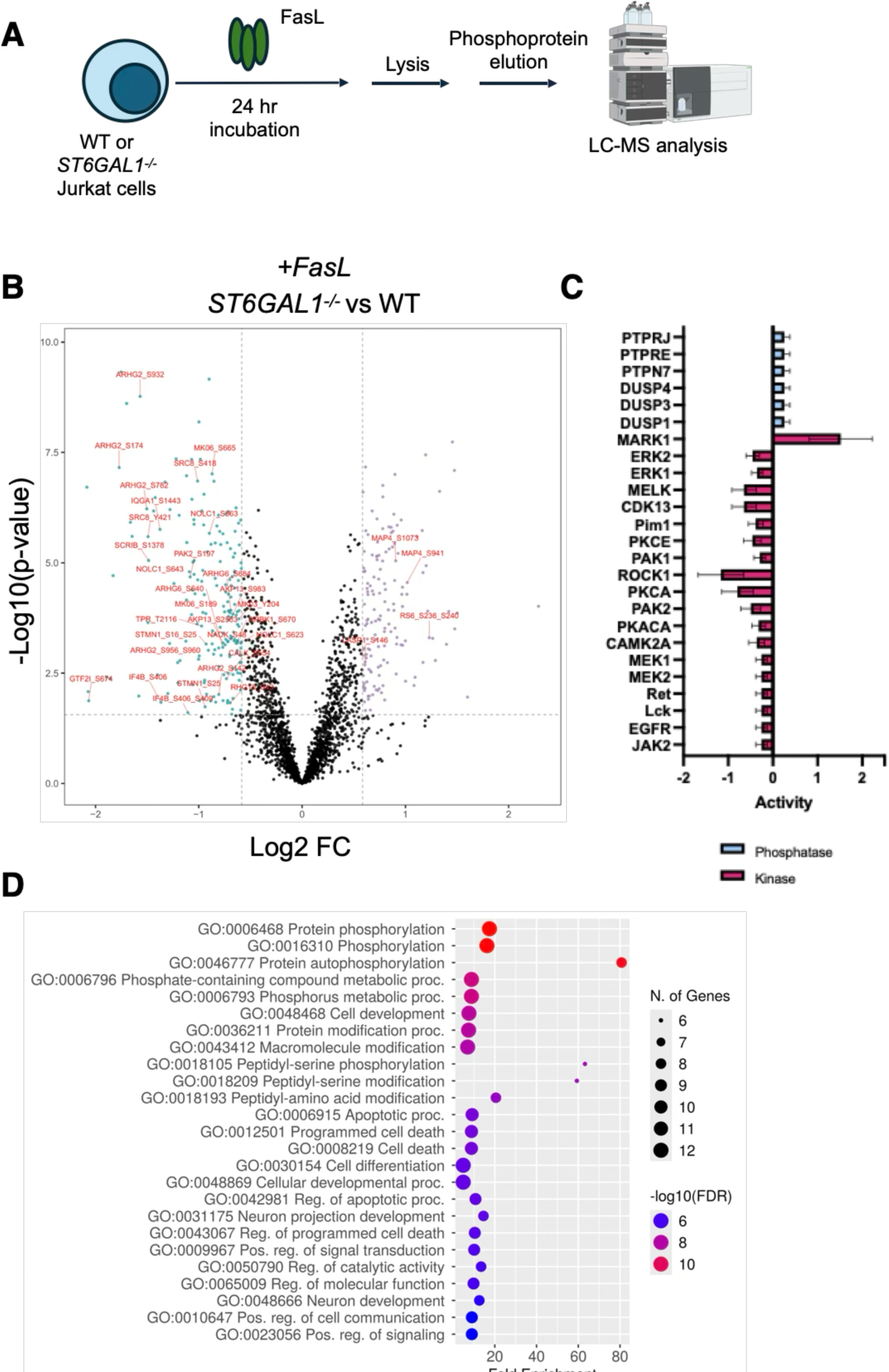
Sialylation influences FasL mediated cell death by altering intracellular phosphorylation events. *A*, Phosphoproteomics workflow for analysis of WT or *ST6GAL1^-/-^*Jurkat cells treated with FasL. *B*, Volcano plot of differentially phosphorylated peptides detected in *ST6GAL1^-/-^*relative to WT Jurkat cells treated with 100 ng/mL FasL. Phosphopeptide IDs that were compatible with RoKAI analysis are annotated in red. Companion plot for cells that were not treated with FasL is presented in Fig. S4. *F*, Differential activity of the kinases and phosphatases in *ST6GAL1^-/-^* relative to WT Jurkat cells identified via RoKAI analysis of the dataset from panel *E*. *G*, GO biological process terms associated with the kinases and phosphatases identified in panel *F*. Figure generated using ShinyGO V. 0.85 (54). All data is the result of four technical replicates submitted for phosphoproteomic analysis.

### Sialic acid-deficient primary human T cells exhibit increased sensitivity to FasL mediated cell death

While Jurkat cells are a useful model system to test fundamental signalling and apoptotic processes, they do not recapitulate all aspects of *bona fide* human T cell immunophysiology. To address this, we next tested if the FasR-FasL cell death axis was sensitive to sialylation in the context of primary human T cells. Here, we isolated T cells from the peripheral blood of healthy adult donors and verified that they expressed FasR via flow cytometry (Figs. 5*A*, *B* and S5). Next, we treated the purified T cells with a recombinant pan-sialidase enzyme from *Vibrio cholerae* (VC). The sialidase efficiently removed sialic acids from α2-6-sialoglycans on T cells as indicated by diminished SNA staining relative to a heat inactivated VC sialidase control group (Fig. 5*D*, *E* and S5). We note that VC sialidase is known to also remove sialic acids linked α2-3 to galactose from primary immune cells since it has broad glycosidase activity against diverse sialosides (16, 35). We next tested the effect of FasL on differently sialylated T cells in multiple ways. First, we executed the Annexin V/DAPI flow cytometry assay on unstimulated T cells and found that both CD4^+^ and CD8^+^ populations were insensitive to FasL treatment regardless of cell surface sialoglycan content (Figs. S6 and S7). We reasoned that FasR expression may not have been high or homogeneous enough on unstimulated cells (Fig. 5B) to measure an effect. To test this possibility, we pre-stimulated T cells *in vitro* using anti-CD3/anti-CD28 beads since it is known that stimulated human T cells express more FasR vs. naïve counterparts (36). Indeed, this approach resulted in a significant increase in FasR expression (Fig. 5B). We then repeated the Annexin V/DAPI assay and found that both stimulated CD4^+^ and CD8^+^ T cell populations were sensitive to FasL-induced cell death, but only following desialylation (Fig. 5*E*, *F*, and S5). The effect manifested as significant decreases and increases in the percentage of live (Annexin V^−^ DAPI^−^) and dead (Annexin V^+^ DAPI^+^) T cells, respectively. We did not observe statistically significant changes across apoptotic populations (Annexin V^+^ DAPI^Int^), suggesting that deislaylation promoted rapid flux through apoptosis to death events. These results support a model where sialoglycans on primary human T cells regulate their sensitivity to FasL-promoted cell death programs.

**Figure 5.**
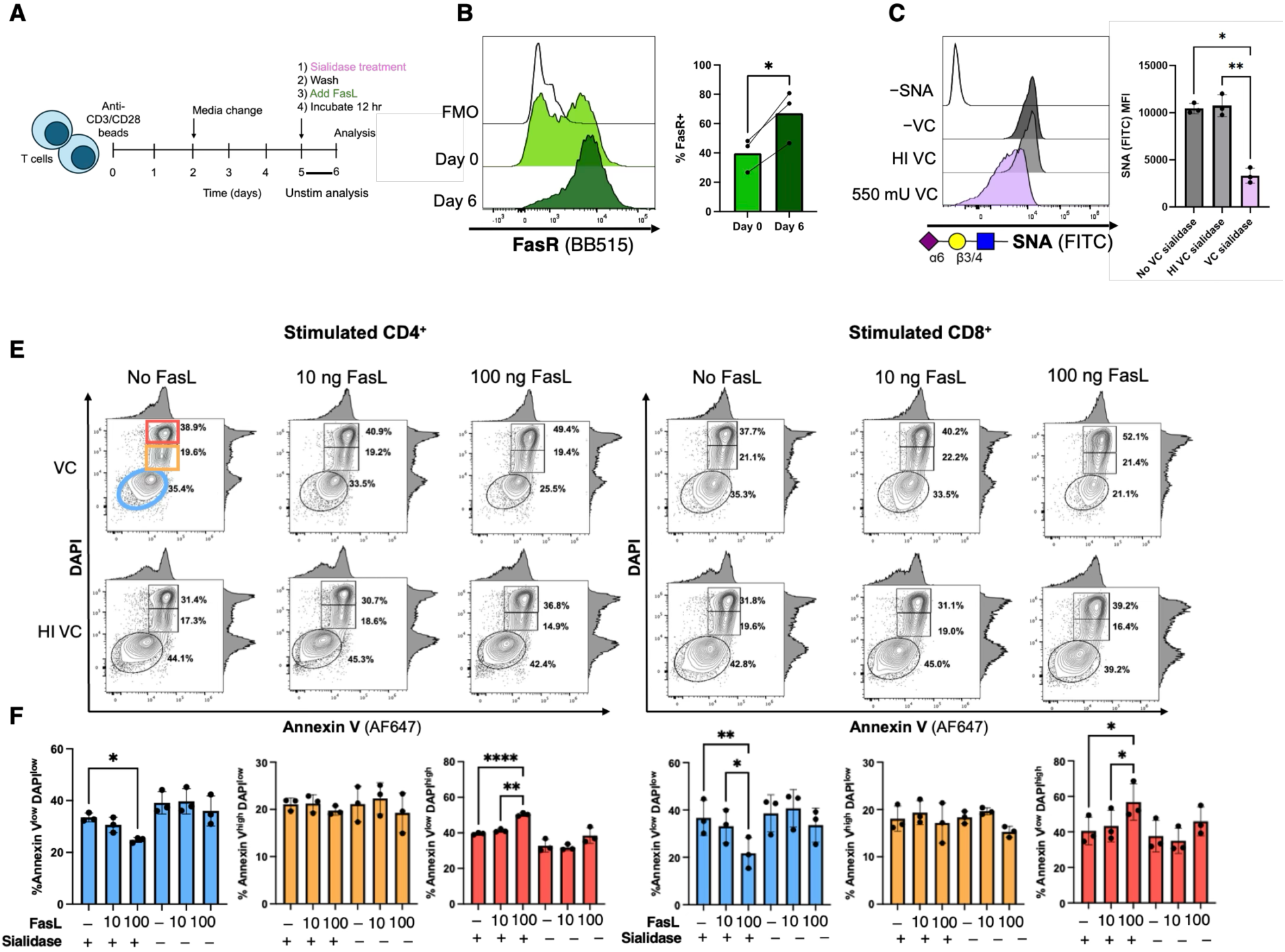
Loss of cell surface sialic acids on stimulated primary human T cells increases cell death in response to FasL. *A*, Workflow for evaluating programmed cell death in primary human T cells following stimulation, desialylation, and exposure to FasL. *B*, FasR expression on unstimulated (day 0) vs. stimulated (day 6) human T cells. *C*, Staining of primary human T cells with SNA for detection of a2-6-sialoglycans following exposure to recombinant VC sialidase (550 mU), heat inactivated (HI) VC sialidase, or no enzyme. *E*, Flow cytometry plots for Annexin V and DAPI staining of stimulated primary human CD4^+^ and CD8^+^ T cells treated with 0, 10, or 100 ng/mL FasL and VC sialidase (550 mU) or HI VC sialidase. Bottom left elliptical gates (blue) = live cells, square gates (orange and red) = apoptotic and dying cells respectively. Percent values indicate the portion of total singlet cell events within each gate. *F*. Quantification of data from *E*. For plots in *B*, *C*, and *F*, results are reported as mean SD from three independent experiments. \**p* < 0.05, \*\**p* < 0.01, \*\*\*\**p* < 0.0001. Student’s t-test, two-tailed (*B*) or one-way ANOVA with Tukey post-hoc test (*C* and *F*). For plots in *E* (and Fig. S7) minor adjustments to the position of gates for the three populations were made to fully capture cell events in the relevant populations due to variation in positioning across different human donors. All gates used for data quantification across all donors are presented in Figs. S8 and S9.

## Discussion

Cell death promoted by the TNF receptor superfamily is essential for maintenance of lymphocyte homeostasis (1, 37). The Fas pathway specifically has been shown to execute complex and opposing pro-apoptotic and pro-survival programs in T cells. For example, it has been shown that T cells can receive pro-survival co-stimulatory signals via FasR-FasL interactions when combined with antigen specific TCR stimulation (38). While this stimulatory role for FasR-FasL dominates at early stages of an immune response, the effect is reversed at later stages when T cells have received repeated TCR stimulation and instead respond to FasL with pro-apoptotic programs (38). This process is important for prevention of T cell-driven autoimmune diseases, such as multiple sclerosis (39). In contrast, FasL-mediated cell death can be detrimental in settings of chimeric antigen receptor (CAR) T cell immunotherapies, since persistence of the CAR T cells is critical for maintaining their anti-cancer effects (40). Recent work has demonstrated that disabling FasR signalling in CAR T cells can improve persistence without compromising CAR-mediated destruction of target cancer cells (41, 42). While all of this previous work underscores essential roles for the Fas pathway in maintaining T cell immune homeostasis, regulatory roles beyond the FasR-FasL protein-protein interaction remain underexplored. The present study begins to address this gap, as we have confirmed that regulation of the Fas pathway in T cells also occurs at the sialoglycan level. Our experiments using unstimulated primary human T cells agree with a non-apoptotic role for the Fas pathway in the context of naïve T cells (36), irrespective of sialoglycan content, whereas stimulated counterparts treated with FasL became maximally apoptotic following treatment with VC sialidase.

These results have important implications when placed into context with our previous work where we found that sialoglycan biosynthesis in human T cells was greatly increased following stimulation through the TCR and CD28 (19). Specifically, stimulated human T cells upregulated *ST6GAL1* and produced dramatically increased amounts of α2-6-sialoglycans as a result. Since our experiments using *ST6GAL1*^-/-^ Jurkat cells confirmed that loss of α2-6-sialoglycans increased susceptibility to cell death programs (Fig. 2*C*, *D*, and *E*), we propose that one role for increased levels of these glycans on primary T cells may be to attenuate Fas pathway signalling. This may help to preserve select antigen-experienced T cells that would be required for production of a memory pool. Scenarios where α2-6-sialoglycans are diminished on activated T cells – for example, following exposure to a sialidase – release this protective suppression and permit more efficient programmed cell death.

While our experiments using Jurkat cells specifically implicated α2-6-sialoglycans as regulators of Fas signalling, companion studies using primary human T cells were less definitive. This was because our enzymatic desialylation approach involved the pan-specific VC sialidase enzyme which cleaves sialic acids from all sialoglycans, including from α2-3-linked structures. While restricted access to specific linkages of sialic acid is a limitation of this approach (α2-6-specific sialidases are lacking), these results are complementary to experiments involving KO of *ST6GAL1*. This is because genetic deletion of ST6GAL1 reduces sialylation at the level of glycan biosynthesis, whereas enzymatic desialylation of a cell surface acts on glycans that have already been produced. Since we observed that both KO of ST6GAL1 and enzymatic desialyation potentiated FasL-promoted cell death, we conclude that effects on the Fas pathway are the direct result of reduced sialoglycan levels at the cell surface as opposed to indirect effects on global glycoprotein biosynthesis and export. Even so, it remains unclear if sialylation of FasR itself is responsible or if other glycoproteins/lipids on the T cell surface play regulatory roles. While others have directly shown that ST6GAL1 KO resulted in complete loss of α2-6-sialoglycans on FasR (11), biophysical measurements should be performed to better understand if FasR-FasL complexation is directly impacted by differential sialylation.

Our results showcase how a specific change in glycocalyx composition can have downstream consequences on intracellular signalling events that manifest in critical immunoregulatory events. Understanding the consequences of disrupting sialoglycans on T cells is of increasing importance since desialylation has recently been identified as a mechanistically orthogonal approach to potentiate T cell functions in settings of cancer immunotherapy (16, 18, 20). Future work in this space should focus on evaluating the consequences of deislaylation on the susceptibility of various T cell populations to Fas pathway-mediated co-stimulation and programmed cell death.

## Experimental procedures

### Cell culture

Jurkat cells were maintained in RPMI 1640 medium (Gibco) supplemented with 10% heat inactivated FBS, 10 mM HEPES, 0.1 mM NEAA, 1 mM sodium pyruvate in a humidified incubator at 37°C with 5% CO_2_.

Cryopreserved peripheral blood mononuclear cells (PBMCs) from 20-30 year old adult humans were purchased from STEMCELL Technologies (#70025.2). Primary human T cells were isolated from these PBMCs using an EasySep Human T cell Isolation Kit (STEMCELL Technologies, #17951). T cells were cultured in RPMI 1640 medium (Gibco, #11875119) supplemented with 100 U/mL soluble IL-2 (Biolegend #589104), 10% heat inactivated FBS, 10 mM HEPES, 0.1 mM NEAA, 1 mM sodium pyruvate, with or without stimulation in a humidified incubator at 37 °C with 5% CO_2_.

### T cell stimulation

Isolated T cells were quantified using flow cytometry, and anti-CD3 anti-CD28 human Dynabeads (Thermo Fisher #11131D) were added into culture at a 1:1 bead:cell ratio. Cells were incubated for 6 days with medium and IL-2 was replenished on day 3. Cells were then stained and analyzed via flow cytometry.

### Primary T cell sialidase treatment

*Vibrio cholerae* (VC) sialidase was expressed in BL21 *E. coli* with a His-tag for purification via Nickel NTA (BioRad, 12009286) fast protein liquid chromatography (FPLC) using a Ni-NTA column. Endotoxin was removed using a Pierce high capacity endotoxin removal spin column (ThermoFisher, #88274). Sialidase enzyme activity was measured using the 4-MU-NANA assay, as reported previously (16). Enzyme treatment was performed in a humidified incubator (37 °C with 5% CO_2_) for 45 minutes in HBSS with 20 mM MgSO_4_ and 3 mM HEPES (pH 7.5). Heat inactivated (HI) enzyme controls were prepared by denaturing the enzymes for 30 minutes at 60°C.

### ST6GAL1^-/-^ Jurkat cell generation

We cloned a previously reported (43) gRNA sequence targeting ST6GAL1 (5’TGTATCCTCAAGCAGCACCC3’) into the pSpCas9(BB)-2A-GFP plasmid (Addgene plasmid # 48138) using Gibson Assembly (44). The plasmid containing the gRNA sequence was transfected into Jurkat cells using Lipofectamine 3000 (Invitrogen) according to manufacturer’s instructions and incubated at 37 °C with 5% CO_2_ for 48 hours. Single GFP+ cells were FACS sorted into a 96-well tissue culture plate and KO colonies were expanded and validated using Sambucus Nigra Lectin (SNA) (Invitrogen, L32479) staining and flow cytometry.

### RT-qPCR

Total RNA was extracted from cells using a Monarch Total RNA miniprep kit (NEB, T2010S) and reverse transcribed into cDNA using a RevertAid first strand cDNA synthesis kit (Thermo Scientific, K1621). We analyzed expression of ST6GAL1 (5’TCCTCTGGGATGCTTGGTAT3’ and 5’ CGTGCAGGCACTATCGAAGA3’), GAPDH (5’AGGTCGGAGTCAACGGATTT and 5’ATGAAGGGGTCATTGATGGCA3’) and 18S (5’ACCCGTTGAACCCCATTCGTGA3’ and 5’GCCTCACTAAACCATCCAATCGG3’) using RT-qPCR (Roche LightCycler 480). PowerUp SYBR green master mix (Applied Biosystems, 3235026) was used for each reaction.

### Immunoblotting

After death ligand treatment, WT and *ST6GAL1^-/-^* Jurkat cells were lysed with RIPA buffer (Thermo Scientific #89900) and prepared in 2x Laemmli SDS sample buffer with beta-mercaptoethanol. Samples were boiled for 5 minutes at 95°C and then resolved by SDS-PAGE. Proteins were transferred to a PVDF membrane and then blocked with 5% skim milk in Tris-buffered saline with 0.1% Tween 20 (TBS-T) for 1 hour at room temperature. Membranes were incubated with primary antibodies against beta actin (1:2000, Cell Signalling Technologies #4970) and cleaved caspase 3 (1:700, Cell Signalling Technologies #9661) overnight in 5% milk TBS-T. The next day, membranes were washed thoroughly with TBS-T and incubated with an anti-rabbit HRP conjugated secondary antibody for 1 hour at room temperature. Signal was developed using SuperSignal West Pico PLUS chemiluminescent substrate (Thermo Scientific #34580) and captured with the ChemiDoc Imaging System (Bio-Rad).

### Soluble death ligand treatment

WT and *ST6GAL1^-/-^* Jurkat cells were plated at a density of 100,000 cells/mL in 12-well tissue culture plates, treated with death ligands (100 ng/mL TNFa, 100 ng/mL FasL, or 50 ng/mL of TRAIL), and incubated for 24 hours at 37 °C with 5% CO_2_.

Human T cells were treated with VC sialidase on day 6 of culture with or without Dynabead stimulation. Cells were incubated with VC sialidase for 45 minutes at 37 °C and then washed with culture medium. Cells were treated with 10 ng/mL and 100 ng/mL of FasL or 10 ng/mL and 100 ng/mL of TNFa and incubated for 12 hours. After 12 hours, Dynabeads were removed from culture using a DynaMag-5 (12303D, ThermoFisher) and cells were washed in FACS buffer (HBSS without Ca^2+^ and Mg^2+^, 1% BSA and 2 mM EDTA) to prepare for flow cytometry staining.

### Annexin V/DAPI staining

Jurkat cells were washed twice with PBS and stained with AF647 conjugated Annexin V (1:250, Thermo Fisher #A23204) in Annexin V binding buffer (10 mM HEPES, 140 mM NaCl, 6.25 mM CaCl_2_, pH 7.4) for 15 minutes. Cells were quenched with DAPI (1:10 000, Thermo Fisher #D1306) in Annexin V binding buffer. Isolated human T cells were washed once with FACS buffer and stained with AF532 conjugated anti-CD3 (1:200, Thermo Fisher #58-0038-42) for 30 minutes in FACS buffer. Cells were then washed twice in Annexin V binding buffer, stained with Annexin V for 15 minutes, and then stained with DAPI at the same dilutions described for Jurkat cells.

Lectin (SNA) stained Jurkat cells and primary human T cells were prepared by washing the cells in FACS buffer and then staining with FITC conjugated SNA (1:2000, Invitrogen #L32479) for 30 minutes on ice. Cells were then washed, resuspended in FACS buffer, and analyzed on a Cytek Aurora (3 lasers, violet (405 nm), blue (488 nm), and red (640 nm)).

### Cell death receptor staining

Jurkat cells were washed twice with PBS and stained with Zombie NIR viability dye (1:1500, BioLegend #423106). Cells were then washed in FACS buffer and stained with either BB515 conjugated anti-Fas (1:100, BD Biosciences #564596), PE conjugated anti-TNFR1 (1:100, BioLegend #369903), or PE-conjugated anti-DR4 (1:100, Invitrogen #12-664-42). For cell death receptor surface staining, total human PBMCs were washed twice with PBS and stained with Zombie NIR viability dye (1:1500). Cells were then washed in FACS buffer, Fc receptors were blocked (1:250, BioLegend #422302), and cells were then stained with Pacific Blue conjugated anti-CD3 (1:100, BioLegend #300417). Finally, cells were stained with either BB515 conjugated anti-Fas or PE conjugated anti-TNFR1 at the same dilutions described for Jurkat cells. Cells were analyzed on a BD FACSymphony A3-II (5 lasers, violet (405 nm), blue (488 nm), red (637 nm), yellow-green (561), and ultraviolet (349)). Data was analyzed on FlowJo software (V10.10.0, BD Biosciences).

### BD S8 (imaging flow cytometer) protocol

*ST6GAL1^-/-^* and WT Jurkat cells were washed in FACS buffer and stained with BB515 conjugated anti-Fas (1:100) after incubating with 100 ng of FasL. They were then washed with Annexin V binding buffer and then stained with AF647 conjugated Annexin V (1:250) in Annexin V binding buffer for 15 minutes. Cells were then stained with DAPI (1:10 000) in Annexin V binding buffer. Cells were analyzed using a BD S8 spectral imaging sorter (5 lasers, violet (405 nm), blue (488 nm), red (637 nm), yellow-green (561), and ultraviolet (349)) (23). Anti-FasR signal (BB515) was quantified on live and apoptotic cells using FlowJo software (V10.10.0, BD Biosciences). Imaging parameters were analyzed using the Cell View Lens FlowJo plugin.

### Phosphoproteomic sample preparation

WT and *ST6GAL1^-/-^* Jurkat cells, either treated with FasL or untreated (biological quadruplicates), were washed twice with PBS. Then, cells were lysed with SDC buffer consisting of 1% sodium deoxycholate, 10 mM tris(2-carboxyethyl)phosphine hydrochloride), 40 mM chloroacetamide, and 100 mM HEPES, pH 8.5, supplemented with protease inhibitor cocktail (Roche, #11836170001). Proteins were digested using a rapid robotics proteomics (R2-P1) workflow followed by enrichment of phosphorylated peptides using R2-P2, as previously described (26, 45) with minor adaptations. The automated workflow was performed using a KingFisher Flex (Thermo Fisher Scientific). In short, a total of 63.75 µg of protein per sample was digested using 25 µL of hydrophilic magnetic carboxylate SpeedBeads (Cytiva, #45152105050250). Four washes were performed in 80% EtOH, and the digested peptides were eluted in 100 mM triethylammonium bicarbonate with sequencing-grade trypsin (1:50 w/w trypsin to protein) after a 3.5-hour incubation at 37 °C, followed by an elution in water. All samples were dried, resuspended in 100 mM TEAB to achieve a final concentration of 2.5 µg/µL, and then peptides were TMT labeled for 1 hour at 20 °C with 400 RPM of shaking by adding 62.5 µg of different TMT 16-plex channels dissolved in 12.5 µL ACN. The TMT reaction was quenched using a final concentration of 0.4% hydroxylamine, after which samples were dried, and subjected to phosphopeptide enrichmentusing Ti-IMAC HP beads (MagReSyn, #MR-THP005). Washes were performed in three solutions: 80% ACN + 5% TFA + 0.1 M glycolic acid, 80% ACN + 1% TFA, and 10% ACN + 0.2% formic acid (FA). Phosphorylated peptides were eluted in 50% ACN + 2.5% ammonium hydroxide, and filtered through a C18 stage tips to avoid carryover of magnetic beads then dried under vacuum before mass spectrometry analysis.

### LC-MS analysis

The peptides were analyzed on an Orbitrap Ascend Tribrid Mass Spectrometer (46) (Thermo Fisher Scientific) coupled to a Vanquish Neo UHPLC (Thermo Fisher Scientific). First, samples were reconstituted in 0.1% FA and loaded on a C18 trap column (300 µm x 5 mm, 5 µm particles, PepMap Neo) using a 300 nL/min flow with buffer A (0.1% FA). Subsequently, samples were linearly eluted using an 85-minute gradient ranging from 2.2% buffer B (99.9% ACN with 0.1% FA) to 28% buffer B on a C18 analytical column (75 µm x 25 cm, 1.7 µm particles, Aurora Ultimate, IonOpticks). The column was then washed by alternating between 99% and 2% B over 5 minutes to bring the total run time to 90 minutes inclusive. Ionization was performed using a nanospray flex source held at +2.0 kV compared to ground, and the ion transfer tube was set to 275 °C. MS1 scans were collected in the Orbitrap from m/z range 400 to 2000 with a normalized AGC target of 100% (4×10^5^ charges), maximum injection time of 123 ms, and a resolution of 60,000 at 200 m/z. Monoisotopic precursor selection was enabled with precursor charge states between 2 to 6 selected for data-dependent MS/MS scans, and dynamic exclusion was applied for 60 s after 1 count. Precursor ions were isolated with a 0.7 m/z quadrupole filter and fragmented by higher-energy collisional dissociation (HCD) at a normalized collision energy (NCE) of 36% with a normalized AGC target of 100% (5×10^4^ charges), maximum injection time of 123 ms, and resolution of 60,000 at m/z 200.

### Phosphoproteomic data analysis

Raw data were converted using msconvert in Proteowizard (47) to mzML format before being searched using FragPipe (23.0) using the ‘TMT-16-phospho’ workflow settings (48). A FASTA file containing all canonical reviewed proteins from the human proteome was downloaded from Uniprot (August 8, 2025) and used as a database with decoys added by Fragpipe. The fixed modifications were set as carbamidomethyl on C [+57.02146 Da], and TMT 16-plex labeling on K and peptide N-terminus [+304.20715 Da], whilst variable modifications were set as oxidation on M [+15.9949 Da], and phosphorylation on S, T, and Y [+79.96633 Da]. Isobaric quantification was performed using Philosopher through TMT Integrator (49) where quantification was performed on the MS2 level with a 20 ppm mass tolerance.

The search results were subsequently subjected to MSStats analyses (50, 51). Intensities were normalized using median normalization, values were imputed using the MSstats in-built accelerated failure model, and only class 1 phosphopeptides were included with a minimum site localization score of 0.75. For subsequent analysis, a fold-change cutoff of 50% was used, and a 5% FDR cutoff was determined using permutation-based FDR correction (52).

To assign upstream kinases to differentially regulated phosphosites, the robust kinase activity inference (RoKAI) tool (27) was used with the default settings and the human reference proteome. RoKAI kinase and kinase target tables were shortlisted (p < 0.05), assigned to significantly changed phosphosites (−1.5 ≥ FC ≥ 1.5, P < 0.05), and selected subsets of these phosphosites were visualized.

## Data availability

The mass spectrometry raw data and search results have been deposited to the ProteomeXchange Consortium via the PRIDE partner repository (53). Datasets used for RoKAI analysis are included in supplementary Table 1.

## Supporting information

This article contains supporting information.

## Acknowledgement

We would like to thank: Dr. Tania Watts for donating the parent Jurkat cell line; Dr. Birinder Ghumman for valuable tissue culture advice; Dr. Feng Zhang for the pSpCas9(BB)-2A-GFP plasmid; Dr. Nathalie Simard and the Temerty Faculty of Medicine Flow Cytometry Core Facility for assistance with S8 operation; David Badillo for reagent titration and optimization; and Dr. Peter McPherson for productive discussions.

## Author contributions

V. A., Q. L., T. S. V., E. S., H. Choksi, F. N. I., Q. S., H. Cui, N. M. R. and L. J. E writing—review & editing; H. Cui, N. M. R. and L. J. E. supervision, funding acquisition, and project administration; V. A. and L. J. E. conceptualization and writing—original draft; V. A., T. S. V., E. S. and L. J. E. data curation, formal analysis, and visualization; V. A., Q. L., T. S. V., E. S., H. Choksi, F. N. I., and Q. S. investigation and methodology.

## Funding and additional information

Research funding was provided to L. J. E. from: The Canadian Institutes of Health Research (CIHR) PTT-190383 & PJT-197877, The New Frontiers in Research Fund NFRFE-2022-00237, The Arthritis Society of Canada IIG-22-0000000126, The Connaught Fund (University of Toronto), The Canada Foundation for Innovation/John R. Evans Leaders Fund, and the Ontario Research Fund: Research Infrastructure. This research was funded in part through a National Institutes of Health Award R00GM147304 (N.M.R), an NIH/NCI Cancer Center Support Grant P30 CA015704 (N.M.R.), and the Searle Scholars Program (N.M.R.). H. Cui acknowledges funding by the Canadian Institutes of Health Research (CIHR) PJT-497271, the Canada Foundation for Innovation/John R. Evans Leaders Fund, and the Ontario Research Fund: Research Infrastructure. V. A. and H. Choksi are grateful for scholarship support from CIHR and The Government of Ontario. Q. L. acknowledges summer research support through the Department of Pharmacology & Toxicology’s Undergraduate Research Opportunity Program (University of Toronto). F. N. I. is grateful for scholarship support from the Indonesian Endowment Fund for Education (LPDP) and the National Research and Innovation Agency of the Republic of Indonesia. Research funding was provided to E. S. by a Washington Research Foundation Postdoctoral Fellowship.

## Abbreviations

The abbreviations used are:

ST6GAL1: ST6 beta-galactosidase alpha-2,6-sialyltransferase 1
WT: wild type
-/-: homozygous knock out
SNA: *Sambucus nigra* lectin
ECL: *Erythina cristagalli* lectin
FMO: fluorescence minus one
FasR: Fas receptor
FasL: Fas ligand
TNFα: tumour necrosis factor alpha
TNFR1: tumour necrosis factor receptor 1
TRAIL-R1: TRAIL receptor 1
TRAIL: TNF-related apoptosis-inducing ligand
Norm.: normalized
MFI: median fluorescence intensity
BB515: BD blue 515
PE: phycoerythrin
DAPI: 4’,6-diamidino-2-phenylin-dole
AF647: Alexa Fluor 647
UT: untreated
CC3: cleaved caspase 3
VC: *Vibrio cholerae*
Log2 FC: log2 fold change
mU: milliunitl
SD: standard deviation.

