## Supplemental Figures for "Sialoglycans on human T cells attenuate death programs executed through the Fas pathway"

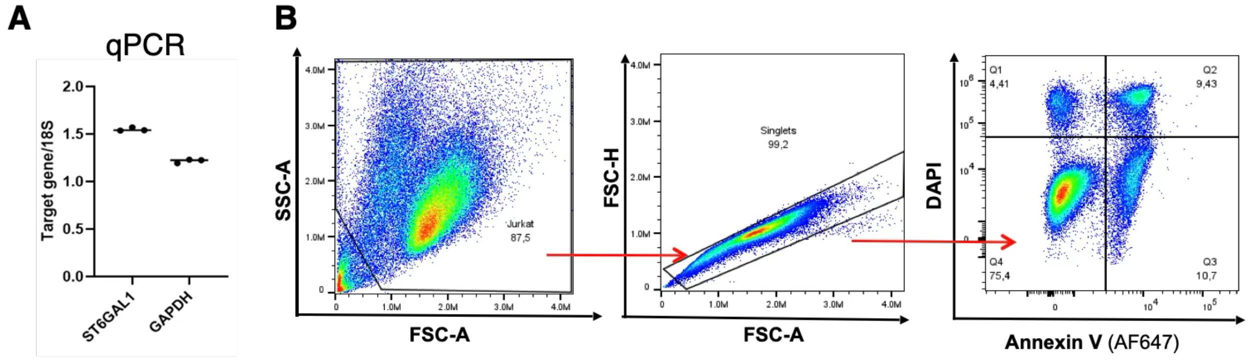


**Figure S1.** *A*, qPCR analysis of ST6GAL1 expression in WT Jurkat cells. Data is reported as normalized to 18S ribosomal RNA. *B*, Representative gating scheme for Jurkat cells treated with 100 ng/mL FasL.

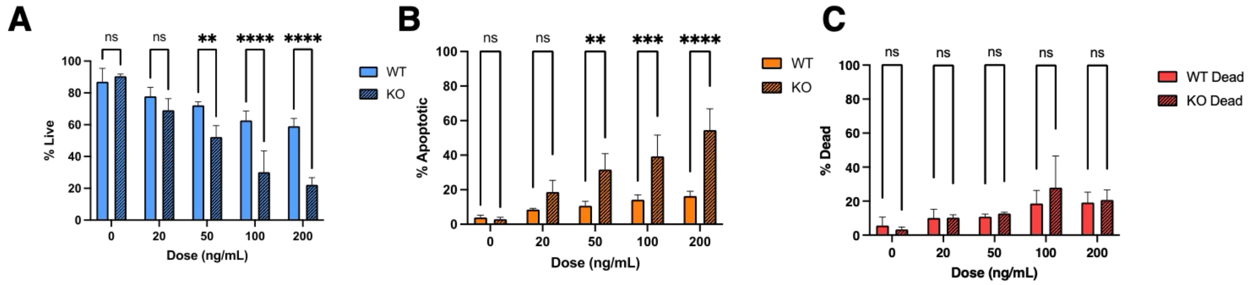


**Figure S2. Dose-response for WT or ST6GAL1$^{-/-}$ Jurkat cells treated with FasL.** *A*, Quantification of live cells. *B*, Quantification of apoptotic cells. *C*, Quantification of dead cells. $^{**}p < 0.01$, $^{****}p < 0.0001$, $p > 0.05$ = ns = not significant. One-way ANOVA with Tukey post-hoc test.

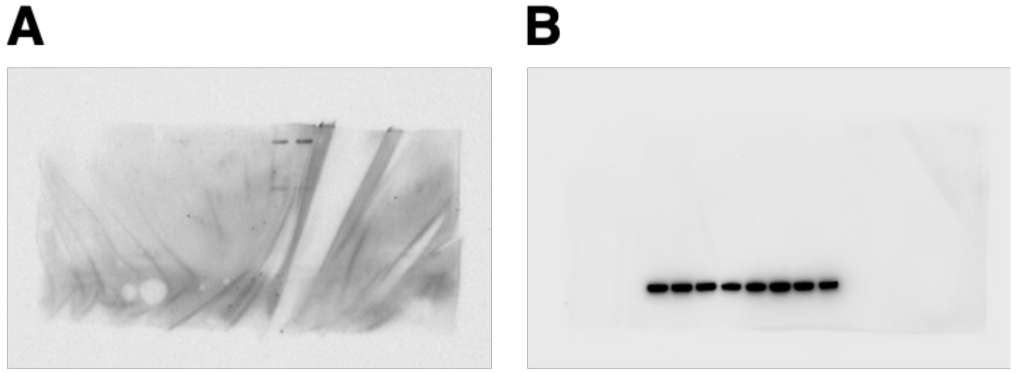


**Figure S3. Uncropped immunoblots.** *A*, detection of cleaved caspase 3. *B*, detection of β-actin. Both panels are companion data to Fig. 2*F*.

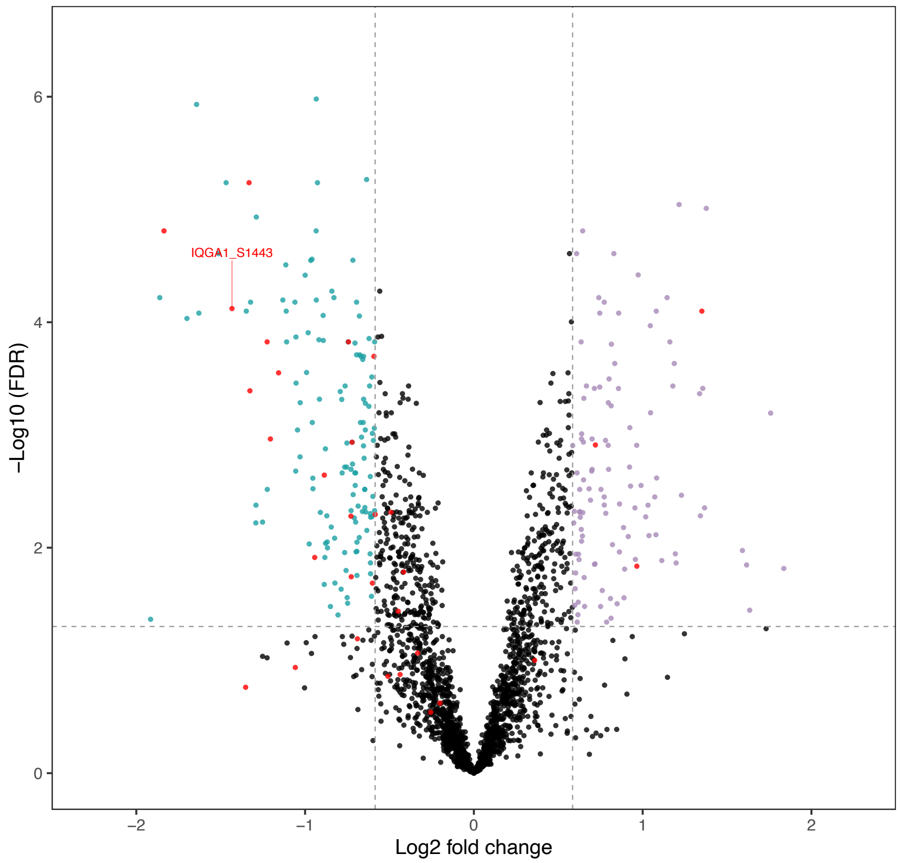


**Figure S4.** Volcano plot of differentially phosphorylated peptides detected in *ST6GAL1*$^{-/-}$ relative to WT Jurkat cells in the absence of FasL. The annotated phosphopeptide was the only statistically significant substrate that could be assigned to kinase activity via RoKAI analysis. Phosphopeptides in red correspond to the substrates identified in Fig. 4B.

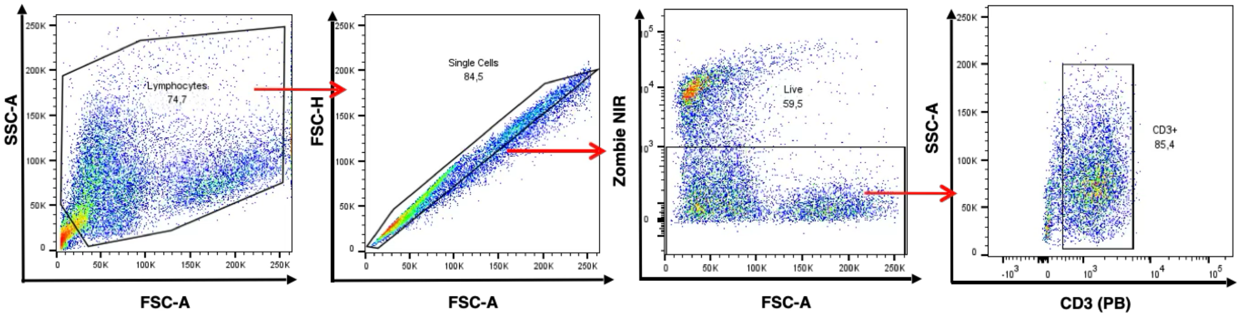


**Figure S5. Gating scheme for evaluating FasR expression and α2-6-sialylation (via SNA) on primary human T cells.** The gating scheme shown is for T cells stimulated (6 days) with dynabeads within a mixture of PBMCs.

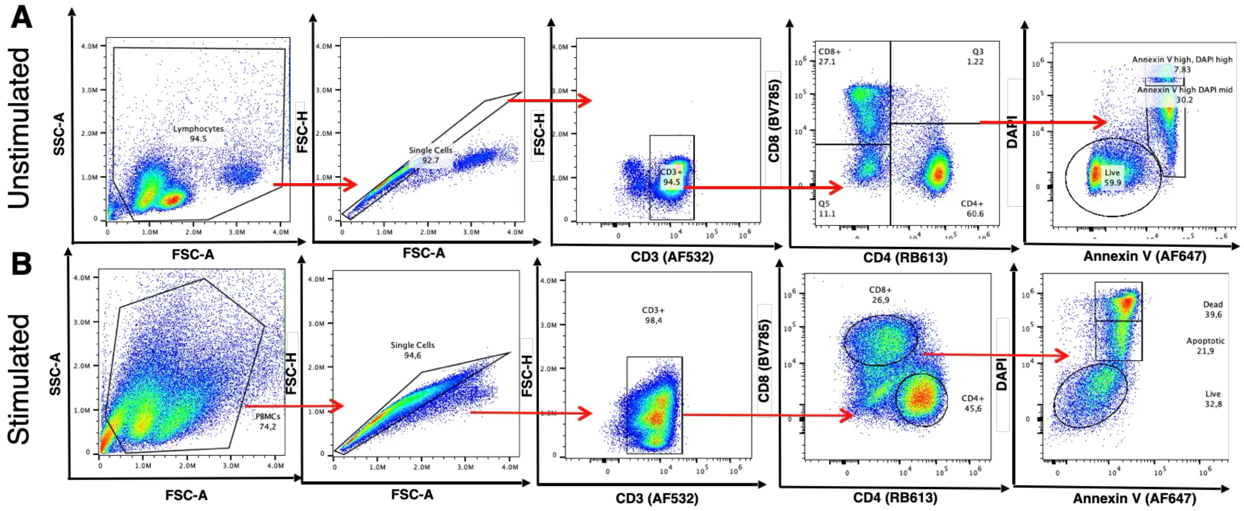


**Figure S6.** Gating schemes for: *A*, unstimulated and *B*, stimulated (dynabeads, 6 days) purified primary human T cells.

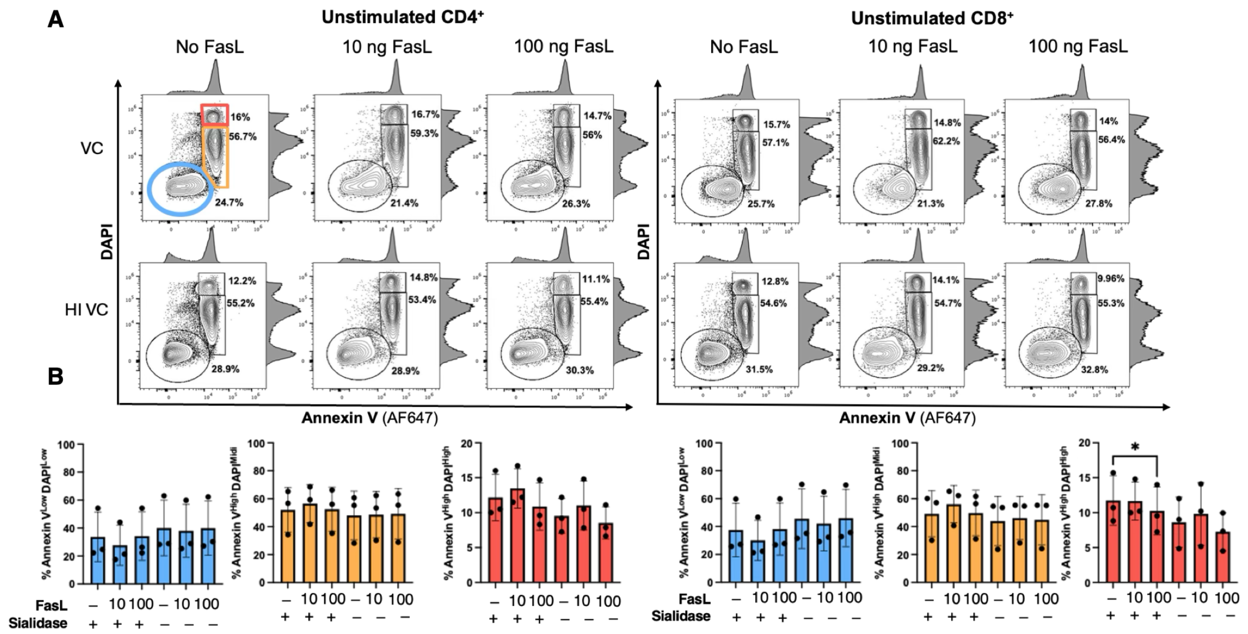


**Figure S7. Sialylation of unstimulated primary human T cells does not influence their sensitivity to FasL mediated programmed cell death.** *A*, Flow cytometry plots for Annexin V and DAPI staining of unstimulated primary human $CD4^+$ and $CD8^+$ T cells treated with 0, 10, or 100 ng/mL FasL and VC sialidase (550 mU) or HI VC sialidase. Bottom left elliptical gates (blue) = live cells, square gates (orange and red) = apoptotic and dying cells respectively. Percent values indicate the portion of total singlet cell events within each gate. *B*. Quantification of data from *A*. Results are reported as mean SD from three independent experiments. **p* < 0.05. One-way ANOVA with Tukey post-hoc test (*C* and *F*). For plots in *A* minor adjustments to the position of gates for the three populations were made to fully capture cell events in the relevant populations due to variation in positioning across different human donors. All gates used for data quantification across all donors are presented in Fig. S7.

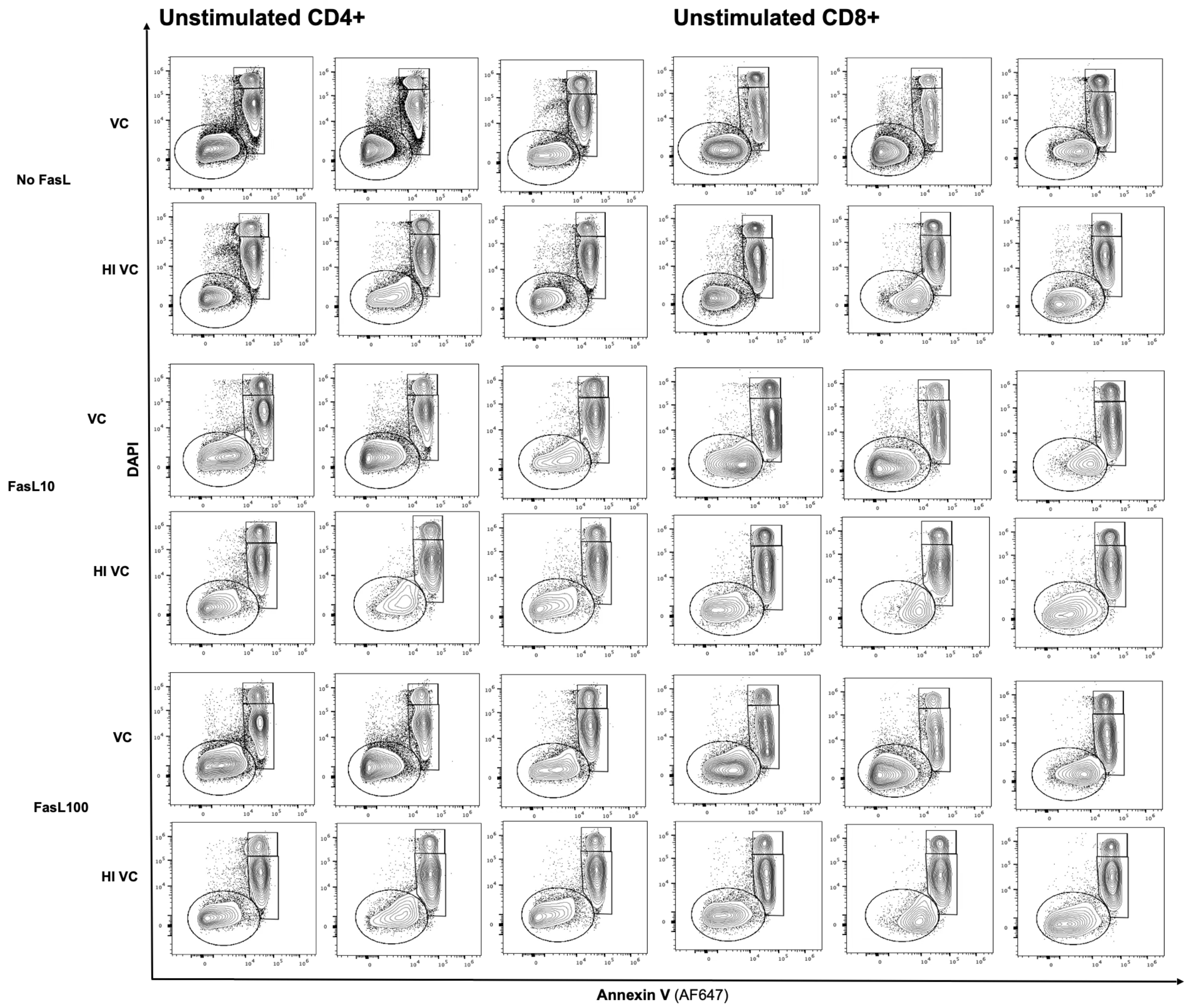


**Figure S8. Gating schemes for live, apoptotic and dead unstimulated T cells across all human donors.**

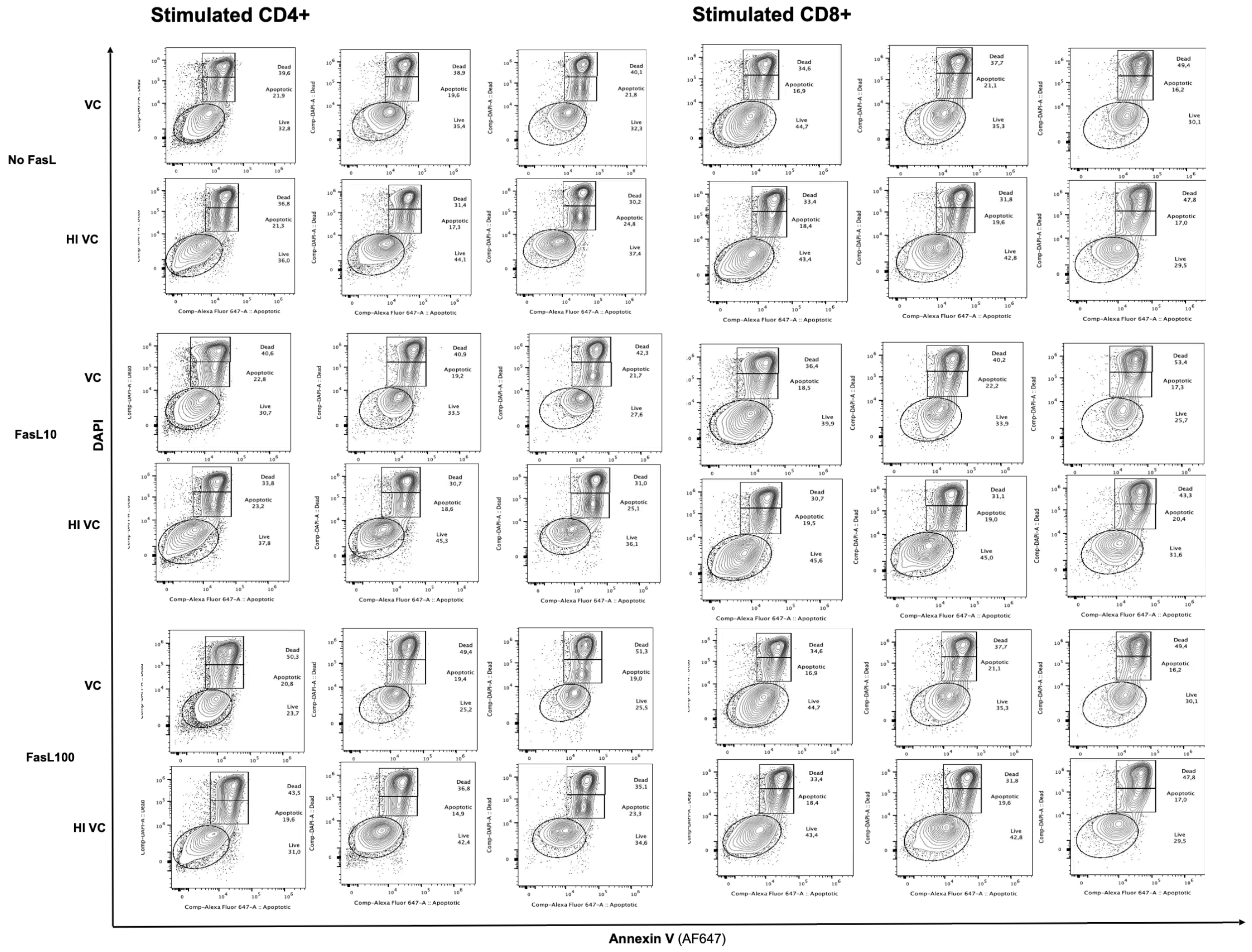


**Figure S9. Gating schemes for live, apoptotic and dead stimulated T cells across all human donors.**
